# Words in context: tracking context-processing during language comprehension using computational language models and MEG

**DOI:** 10.1101/2020.06.19.161190

**Authors:** Alessandro Lopopolo, Jan Mathijs Schoffelen, Antal van den Bosch, Roel M. Willems

## Abstract

The meaning of a word depends on its lexical semantics and on the context in which it is embedded. At the basis of this lays the distinction between lexical retrieval and integration, two basic operations supporting language comprehension. In this paper, we investigate how lexical retrieval and integration are implemented in the brain by comparing MEG activity to word representations generated by computational language models. We test both non-contextualized embeddings, representing words independently from their context, and contextualized embeddings, which instead integrate contextual information in their representations. Using representational similarity analysis over cortical regions and over time, we observed that brain activity in the left anterior temporal pole and inferior frontal regions shows higher similarity with contextualized word embeddings compared to non-contextualized embeddings, between 300 and 500 ms after word presentation. On the other hand, non-contextualized word embeddings show higher similarity with brain activity in the left lateral and anterior temporal lobe at earlier latencies – areas and latencies related to lexical retrieval. Our results highlight how lexical retrieval and context integration can be tracked in the brain using word embeddings obtained with computational models. These results also suggest that the distinction between lexical retrieval and integration might be framed in terms of context-independent and contextualized representations.

## Introduction

Sentence comprehension is thought to rely on two basic operations, lexical retrieval and integration [1–6]. Lexical retrieval refers to the process that extracts the meaning of individual words from the mental lexicon [1, 2, 6], which takes place before an integration process combines the individual semantic representations in meaningful ways.

Baggio & Hagoort [7] argue that it is difficult to directly test the nature of the distinction between retrieval and integration using traditional task-based experimental paradigms, since it might be impossible to devise a task tackling only one operation while leaving the other untouched. Therefore, in the study presented here, we instead propose to use computational semantic models to further investigate the neural basis of these two basic operations.

In the present study, we use computational semantic models to investigate the neural basis of lexical retrieval and integration. We use two classes of computational semantic models, representing linguistic items either as independent or dependent from their context of occurrence. In the computational linguistics literature, these models are usually referred to as non-contextualized and contextualized embeddings, respectively [8, 9]. By comparing them to MEG data collected during sentence comprehension, we aim to get more insight into the neural basis of these processes by showing that integration is approximated by contextualized embeddings and that it is a separate process (both functionally and physiologically) from semantic memory. The main methodological contribution of computational modeling to the study of language processing in the brain relies on the fact that it helps in avoiding the limitations of task-oriented studies by exploiting the richness of naturally occurring sentences and by relying on a fleshed out model of the investigated processes. In other words, computational linguistic modeling provides a more direct implementation of the process and does not require the assumption that the process can be decomposed in orthogonal subprocesses that can be controlled by specific experimental tasks.

### Integration as contextualization

The distinction between retrieval and integration can be grounded on the observation that the language system seems able to deal with a virtually infinite number of utterances, which in turn seem to be composed by a limited, although flexible, set of primitives, such as phonemes, morphemes, words, and phrases. By storing these primitives in a hypothetical partition of long-term memory, recalling them when necessary, and combining them in a seemingly unbound number of configurations, humans can deal with a large variety of messages in a parsimonious, flexible and creative way.

The present study frames the concept of integration in terms of contextualization of lexical items. This is based on the assumption that the meaning of a word depends on how it is represented in memory, and how this representation changes as a result of its integration in context, linguistic in our case, it is presented in.

1. In order to open a new account, you should go to a ***bank***.
2. A fisherman is sitting with his rod on the ***bank*** of the river Thames. For instance, humans distinguish the meaning of *bank* as “building or financial institution” or as “the shore of a river” depending on whether it is encountered in the context of Sentence 1 or 2. The presence of the string *“a new account”* in Sentence 1 steers the interpretation towards the financial domain, whereas the string *“a fisherman”* acts as bias towards a river-related interpretation of the word ***bank***. A distinct yet similar disambiguation problem arises with tokens that are not lexically ambiguous, such as *dog* in Sentences 3 and 4. Although the basic meaning of the term is the same in the two sentences, the two tokens take on two distinct sentential roles: as the object of an action (4) or the subject of a statement realized as a compound nominal predicate (3).
3. The domestic ***dog*** is a member of the genus Canis, which forms part of the wolf-like canids.
4. I took my ***dog*** out for a walk in the park.

It has been suggested that the human brain creates representations of words that are different according to such contextual cues [10].

Recent years have seen an increase of interest in the role of context in the construction of artificial representations of word meanings in the field of computational linguistics. Striking results have been reported with so-called contextualized word embedding models, showing significant improvements in performance in several computational linguistic tasks as compared to those obtained using word embeddings (representations) that do not consider context [9, 11–14]. In this study, we investigated the effect of contextualization of words on brain activity, comparing non-contextualized and contextualized models. Our working hypothesis is that, after lexical retrieval from memory, brain activity can be modelled by non-contextualized embeddings. Conversely, the product of lexical integration can be modelled by contextualized embeddings.

Brain activity related to language comprehension reflects processes that involve different areas of the brain at different moments in time following the onset of the stimulus [15–17]. It is therefore capital to show that the putative similarity between a model and a brain process concerns not only areas associated with such process, but also that it does so in a time frame that is compatible with the time course of language processing. For this reason, we use a magnetoencephalographic (MEG) dataset collected during sentence reading. MEG records brain activity at the level of milliseconds, and with a reasonable anatomical resolution, making it ideal for a study interested also in the when, and not only the where of a specific neural process [18].

## Lexical processing in the brain

### Neural loci

**Semantic memory**, a component of long-term memory, acts as the storage of knowledge and representations of basic linguistic units, such as words. In a simplified manner, we can define memory as the mental lexicon, or the equivalent of a vocabulary in which the representations of words are stored and wait to be retrieved during production or comprehension. Binder and colleagues [19, 20] provide a comprehensive picture of the cortical areas substantiating semantic memory. Memory, semantic memory in particular, is associated with the lateral portion of the left temporal cortex (middle temporal gyrus), parts of the inferior temporal gyrus, and the inferior parietal cortex. An important role is also hypothesized for the anterior portions of the temporal lobe (anterior temporal pole, ATP). The involvement of the ATP is confirmed by both studies on semantic dementia [21, 22], and by a large body of neuroimaging literature [23–26]. These findings have been summarized by Patterson & al. 2007 [27] and led to the formulation of the hub and spoke model, which posits that concepts are represented by a network of sensorimotor representations converging in the ATP, which acts as a hub collecting and controlling modality-specific features in order to produce supra-modal representations.

**Integration** is a process that operates on representations retrieved from semantic memory. In its most basic formulation, integration consists of merging two linguistic tokens (e.g., two words) and creating a larger unit, such as a phrase or, more simply, a bi-gram. Integration is an operation that takes a token and embeds it into the context represented, for instance, by the other tokens making up the sentence in which it is presented. Brain imaging and brain lesion studies suggest that the inferior frontal gyrus, in interaction with areas in the perisylvian and temporal cortex, plays an essential role in lexical integration [17, 28].

Integration also involves anterior temporal areas. By contrasting the activity recorded during the reading of sentences and of word lists, works such as Mazoyer & al. (1993) [29], Stowe & al. (1998) [30], Friederici & al. (2000) [15], Humphries & al. (2006) [31], and Humphries & al. (2007) [32] reported an increase in activity in ATP for the former condition as compared to the latter. The role of ATP in processing integration is confirmed by another series of studies narrowing down the scope of the analysis. Rather than working with sentences as a whole, these analyses focused on simple phrasal processing, consisting of the composition of a wide range of phrasal and syntactic compositional types and cross-language and modality [33–37].

### Timing of processes

Besides the cortical loci of processing, sentence processing is characterized by a specific temporal profile that describes the timing of each of its sub-processes [16, 32]. The earlier stages mainly concern the recognition of the word from its auditory (for spoken words) or graphic (for written words) form and involve primary auditory or visual areas between the onset of a word and 150–200 ms. The phases that interest our analysis are the so-called Phase 1 and Phase 2, as described by Friederici [16].

**Phase 1** takes places after the word form has been identified, and can be broken down into sub-phases. First, around 200 ms after the onset of a word, its category is identified (i.e., whether it is likely to be a verb, a noun, an article, etc.). Subsequently, information of its lexical meaning is retrieved from semantic memory implemented in the middle temporal gyrus. This process takes place approximately between 150 and 300 ms after the onset of a word.

**Phase 2** follows between 300 and 500 ms after stimulus onset. It roughly corresponds to integration, as introduced in the previous Section. In this phase, the lexical representation of a word retrieved in Phase 1 is embedded in the contextual representation consisting of the retrieved and unified representation of the other words composing the sentence that is processed [38–41].

## Computational models

For the purpose of this study, we use two types of computational models developed for generating word representations: non-contextualized models and contextualized models. These models create so-called word embeddings, which consist of vectors of real numbers populating a high-dimensional space. In other words, a model *M* takes a word *w* and returns a real vector 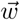 representing *w* in a high-dimensional space *S*.

The first type of model generates representations 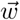 that are independent from the context (sentence, paragraph, etc.) in which the represented word *w* is located. We call this type of model non-contextualized, and it is represented by the popular word2vec model (Section 4) [8].

Besides non-contextualized models, we also consider a contextualized model: ELMo (Sections 4 [9]. This model, contrary to word2vec, assigns representations 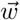 that depend on the textual context in which the represented word *w* is located. For instance, if the word *dog* appearing in Sentences 3 and 4 always obtains the same embedding 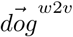 from word2vec, it will obtain two different vectors 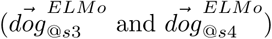, one for each of the two contexts in which it is found.

As shown in Figure 1, this becomes evident when we compute the similarity between the embeddings. The cosine similarity between the word2vec generated 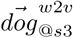 vector (Sentence 3) and 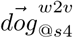 (Sentence 4) is 1.0, indicating a total identity between the two representations. Instead the cosine similarity between the ELMo generated 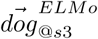 vector (Sentence 3) and 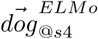 (Sentence 4) is 0.92. Similarly the similarity between the word2vec representations of *bank* (Sentences 1 and 2) is 1.0, whereas the similarity between the ELMo representation of the same words is 0.87.

**Fig 1.**
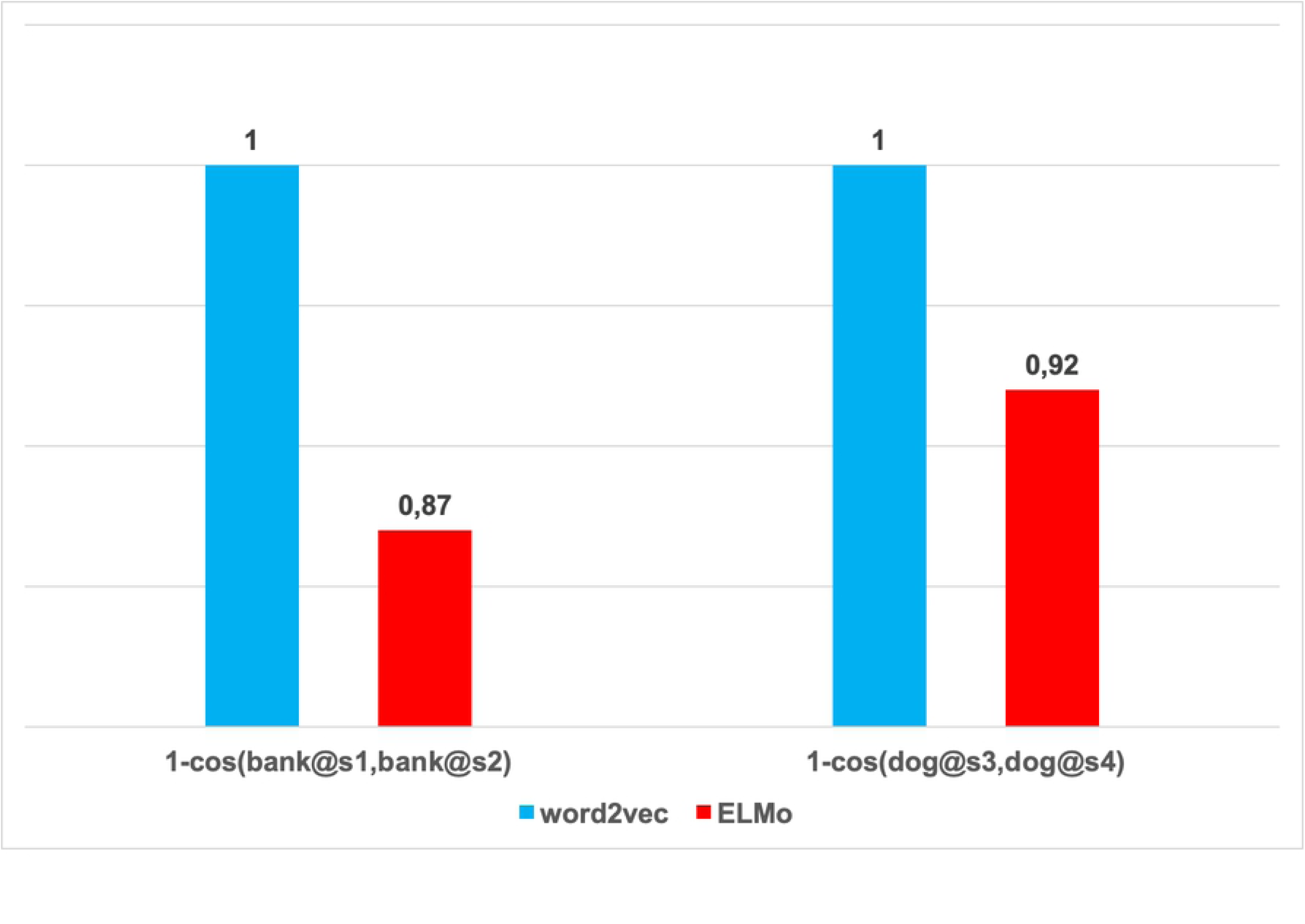
The effect of contextualization on pairwise word similarity. Word2vec (blue) returns identical representations for words independently from the sentences they are presented in. This is evident from the 1.0 cosine similarity between *bank* in Sentences 1 and 2 – on the one hand – and between *dog* in Sentences 3 and 4. Conversely ELMo (red) instead produces different contextualized representations of the same words depending on the context they are used, note the lower similarities (0.92 and 0.87).

### Non-contextualized embeddings (Word2vec)

Word2vec is an artificial neural network-based model used to produce word embeddings. It has been proposed as a more advanced alternative to earlier distributional sem ntic vector spaces such as latent semantic analysis. From an architectural point of view, word2vec consists of a shallow, two-layer neural network. The model can be trained either to predict the current word from a window of surrounding context words (continuous bag-of-words, CBOW), or – conversely – predict the surrounding window of context words given a target word (continuous skip-gram, CSG).

Training creates a high-dimensional vector space populated by word vectors, which are positioned in the space in such a way that words that share similar semantic and syntactic properties lay close to one another. Fig 2 represents the architecture of the CBOW version of word2vec which is used in the present study (adapted from [42]). The trained embeddings correspond to the weights stored in matrix *W*, whose dimensions [*v × d*] correspond to the size of the modeled vocabulary (*v*) and the chosen number of dimensions (*d*) of the vector space itself. Therefore, the way word2vec assigns embeddings to word *w* can be seen as a sort of dictionary “look-up” where the embedding of word *w*_*i*_ corresponds to row *i* of matrix *W*.

**Fig 2.**
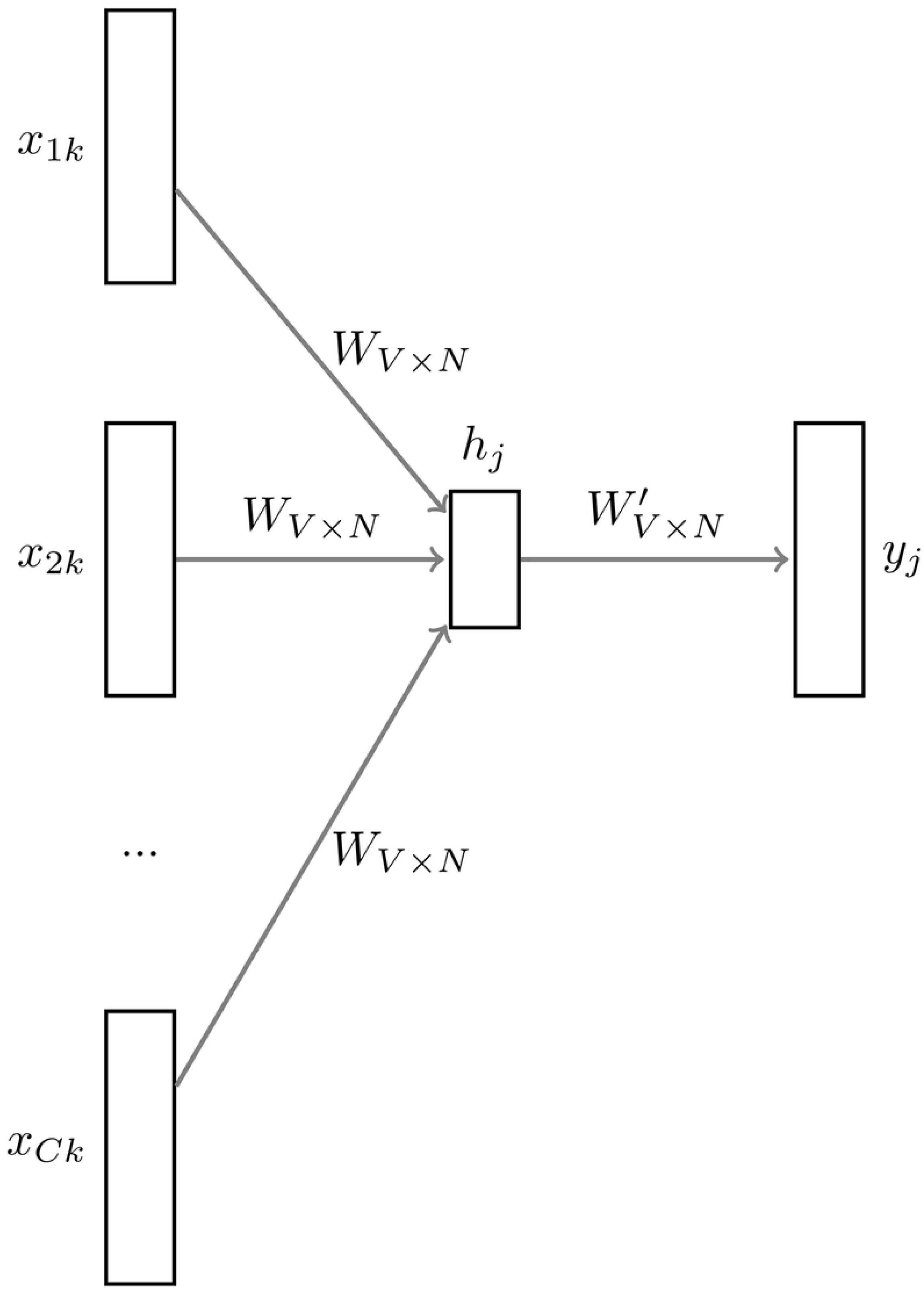
Architecture of the non-contextualized word embedding model (word2vec). The model embeddigns (*W*) are trained by predicting the current word from a window of surrounding context words (continuous bag-of-words, CBOW).

For the purpose of the present study, it is important to point out the role of context with regard to the way word2vec is trained and used to assign word embeddings. Context indeed plays a capital role during the training of the model. In both cases, the context of a word is present in the pipeline, either as the input or as the target of the training function. Nonetheless, once the model is trained, its application is blind to the context and relations that the words have.

### Contextualized word embeddings (ELMo)

The contextualized word embedding model ELMo [9] relies on the properties of recurrent neural networks. Contrary to word2vec, ELMo is a deep contextualized model for the generation of the word representation. It models complex characteristics of word use that vary across linguistic contexts.

From an architectural point of view, ELMo is a recurrent bi-directional language model (biLM) composed of 2 layers of bi-directional recurrent units (implemented as LSTM layers) feeding to an output softmax layer. A language model refers to a system (stochastic or neural) trained on predicting a word given its preceding context. In its most common formulation, a language model is an approximation of the function *P* (*w*_*i*_*|w*_1:*i−*1_), i.e. a function that assigned a probability to a word *w*_*i*_ given its prefix *w*_1:*i−*1_.

A recurrent layer is a layer that creates a representation of the input sequence (sentence) at word *w*_*i*_ as a combination of the representation of the *i*_*th*_ word and the representation of its preceding (if the recurrent neural network proceeds left to right) or following (if it proceeds right to left) context.

Fig 3 illustrates the structure of the ELMo model consisting of a 2-layer biLM. ELMo’s bidirectionality refers to the fact that its hidden layers receive information about both the preceding and following words of *w*_*i*_. The representation ELMo generates for a word consists of a concatenation of the activation of its two recurrent layers.

**Fig 3.**
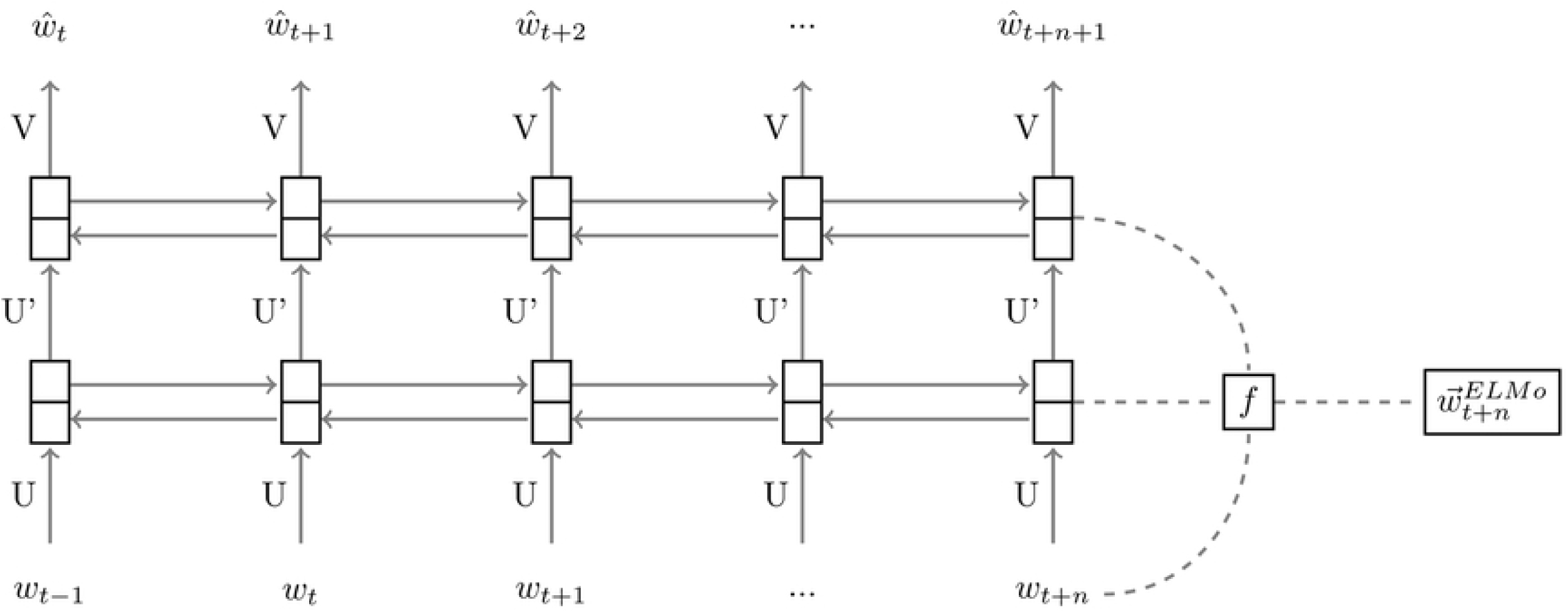
Architecture of the contextualized word embedding model (ELMo). ELMo architecture consisting of a 2-layer biLM, where the layers are implemented as LSTMs. The contextualized embedding is produced as a linear combination of its components.

Contrary to how word2vec assigns embeddings to words, ELMo does not employ a word-vector dictionary (as word2vec *W* in Fig 2). Given *w*_*i*_ in a sentence *S* = [*w*_1_, *w*_2_, *…w*_*n*_] ELMo instead creates the embedding 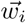 by passing the whole *S* text through the biLM. Embedding 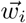 will depend on context *w*_1:*i−*1_ and *w*_*i*+1:*n*_ as the combination of the activation of the LSTM layers corresponding to the presentation of *w*_*i*_ to the model. For these reasons, if *w*_*i*_ appears in a different sentential context *S*′, its 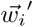 will be different.

### Training

Since the MEG data (described in Section 4) was collected from Dutch native speakers reading Dutch sentences, we used models trained on Dutch texts. Word2Vec was trained on the CBOW task on the Corpus of Spoken Dutch (CGN2). For ELMo we used the pretrained Dutch model provided by the ELMoForManyLangs collection^1^ with the same hyperparameter settings as [14] and trained on a set of 20-million-words Dutch corpus randomly sampled from the raw text released by the CoNLL 2018 shared task on Universal Dependencies Parsing.

### Hypotheses

This study aims at investigating the neurobiological underpinnings of the distinction between non-contextualized and contextual representation as derived from recent developments in word embedding modeling techniques. More specifically, for the **non-contextual model**, we hypothesize a similarity between this model and the operation of **lexical retrieval** from memory. We expect that a model with such characteristics should most closely resemble the activity in lateral temporal regions during processing, around 200 ms after word onset.

Conversely, **contextual model** representations of words are expected to resemble more the activity related to **integration** and described as pertaining inferior frontal and antero-temporal regions between 300 and 500 ms after the onset of the word. Fig 4 schematically represents these hypotheses.

**Fig 4.**
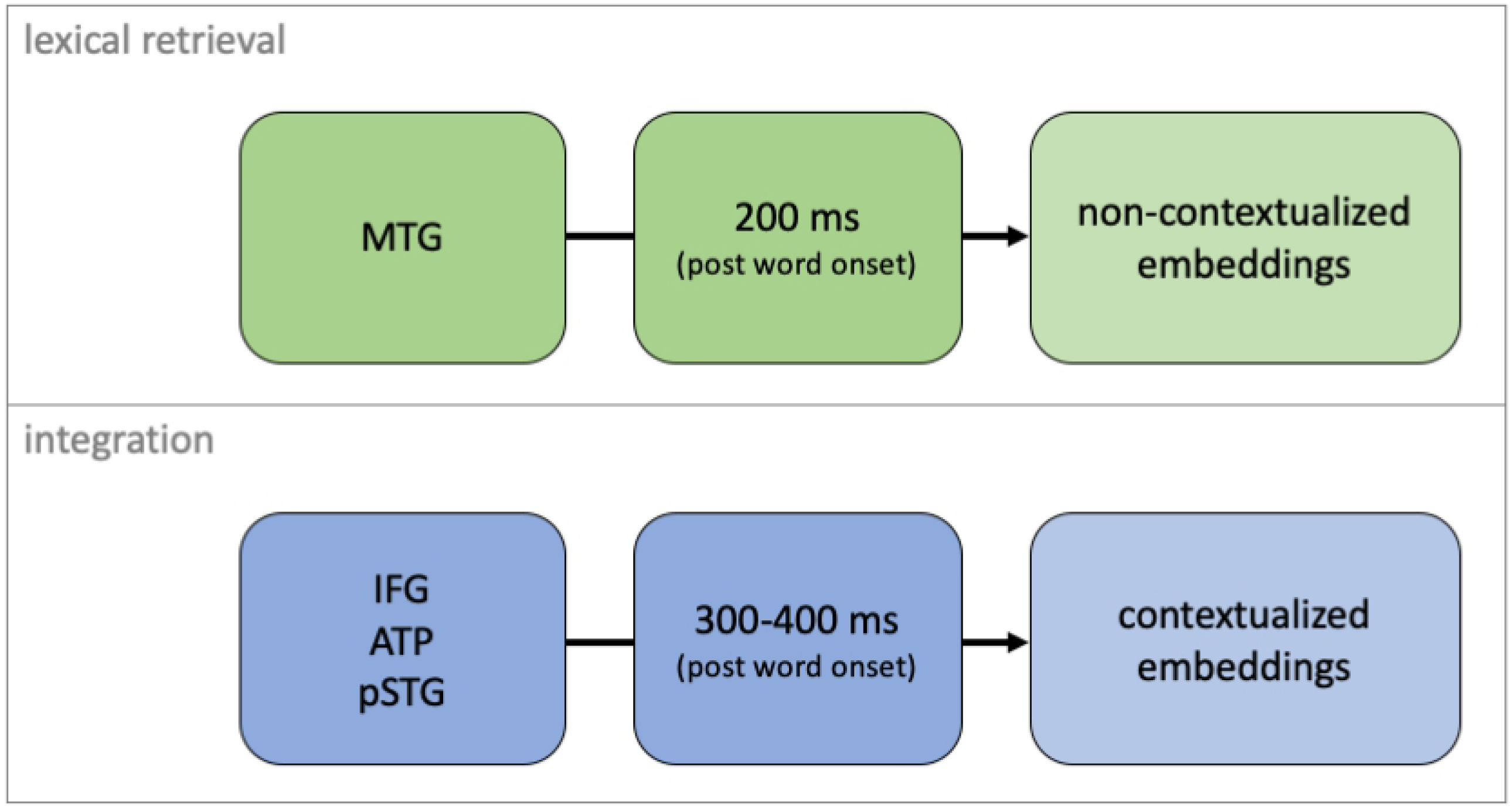
Hypotheses. The hypothesis presented in this paper is that brain activity relative to lexical retrieval can be modelled by non-contextualized embeddings, whereas that of integration can be modelled by contextualized embeddings instead.

## MEG Data

We used the MEG data belonging to the MOUS dataset [18] collected at the Donders Centre for Cognitive Neuroimaging in Nijmegen, The Netherlands. For more details on the acquisition procedure, stimuli, pre-processing, and source reconstruction techniques, we refer to the original paper and to Schoffelen & al. 2018 [18, 45].

### MEG data acquisition and pre-processing

The data were collected with a 275 axial gradiometer system (CTF). The signals were digitized at a sampling frequency of 1200 Hz (the cutoff frequency of the analog anti-aliasing low pass filter was 300 Hz). Head position with regards to the sensors was determined by 3 coils attached to the participant’s head. Electro-cardiogram and the horizontal and vertical electro-oculogram were measured by 3 bipolar Ag/AgCl electrode pairs.

Electrocardiogram artifacts were identified based on their topography and subtracted from the data. The data was segmented into trials corresponding to activity recorded from -183 ms before word presentation to a variable time after word presentation, depending on word length. Trials that contained artifacts (Eye movements and muscle contractions and jump artifacts in the SQUIDs) were excluded from further analysis. Next, the power line interference was estimated and subtracted from the data. The data were down-sampled to a sampling frequency of 300 Hz.

Source reconstruction was performed using a linearly constrained minimum variance beamformer (LCMV) [46], estimating a spatial filter at 8,196 locations of the subject-specific reconstructed midcortical surface. The dimensionality of the data was reduced by applying an atlas-based parcellation scheme based on a refined parcellation Conte69 atlas (191 parcels per hemisphere). After that, spatial filters were concatenated across vertices comprising a parcel. The first two spatial components were selected for each parcel. For more details on this procedure we refer to Schoffelen & al. 2018 [45].

### Subjects

We used the data of 74 subjects belonging to the MEG section of the MOUS dataset [18]. All subjects were Dutch native speakers, who were asked to silently read 120 Dutch sentences presented on a screen word by word, containing a total of 1377 words. All sentences varied between 9 and 15 words in length.

### Stimulation paradigm

The sentences were presented visually with an LCD projector, with a vertical refresh rate of 60Hz situated outside the MEG scanning room, and projected via mirrors onto the screen inside the measurement room. All stimuli were presented in a black mono-spaced font on a gray background at the center of the screen within a visual angle of 4 degrees. Sentences were presented word-by-word with a mean duration of 351 ms for each word (minimum of 300ms and maximum of 1400ms, depending on word length). The duration of the stimuli was determined taking into account both the word length, the number of words in the whole sentence, and the number of letters within each word. Each word was separated by an empty screen for 300 ms before the onset of the next word.

## Analysis

The computational models introduced in Section 4 are used to generate vectorial representations of the same stimulus sentences presented during the acquisition of the MEG data described in Section 4. These representations are generated at the word level, meaning that our models assign a set of real number vectors for each word of the stimulus sentences. The non-contextualized word embedding (word2vec) assigns only one vector per word, whereas the contextualized models (ELMo) represent every word with a set of vectors, each one corresponding with one component (layer) of their internal architecture. In our analyses, a word represented by ELMo is assigned one vectorial representation corresponding to the average of the activation of the two bidirectional layers composing the ELMo network 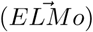.

This section describes the analysis methodologies employed to map the two vectorial representations 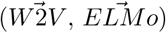 to the corresponding brain activity recorded with MEG. Our goal is to both map the overlap between model representations and brain activity at the anatomical level, and to track the temporal evolution of such similarity. For this reason, we adopted a version of representational similarity analysis (RSA) implemented in a way to return both spatially (anatomical) and temporally situated similarity scores.

### Representational similarity analysis through time

Given a set of linguistic units, for instance, words *w*_1_, *…w*_*n*_, we can generate a vectorial representation for each of them using the embedding models described in Section 4. Words *w*_1_, *…w*_*n*_ are assigned vectors 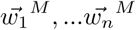 from the computational model *M*. These are drawn from a representational space *RS*^*M*^ populated by the vectorial embeddings. At the same time, it is possible to derive a brain representation of the same words. This representation is the word-wise single trial activity recorded in the MEG dataset described in Section 4. Note that we did not average over time per trial, but that we kept the signal for each word as is. These activation samples also create a representational space *RS*^*B*^ populated by the same words *w*_1_, *…w*_*n*_. At this point, the aim of the analysis is simply to measure how similar *RS*^*M*^ and *RS*^*B*^ are. In order to do so, we employ representational similarity analysis (RSA). Instead of directly mapping the two spaces, RSA compares pairwise similarities across different spaces.

We conducted RSA at the level of anatomical regions and using a sliding, partially overlapping, temporal window (width 30 ms, step 16 ms). In this way, we obtained a neural representation for each word, for each brain region and time window, which could be paired with a specific model-derived vectorial representation of the same word.

Given a model *RS*^*M*^, we first obtained a representational similarity structure *ss*^*M*^ consisting of a [*n × n*] matrix where element 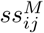 denotes the dissimilarity (computed as Euclidean distance) between embeddings 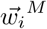 and 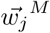 from *RS*^*M*^ of words *w*_*i*_ and *w*_*j*_. Similarly it is possible to derive a series of representational structures 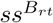 for each anatomical region *r* and time window *t*. Element 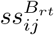 quantifies the similarity between the brain activity in *r* and time *t* for word stimuli *w*_*i*_ and *w*_*j*_. Fig 5 displays the representational similarity structure for brain activity in region *r* and time *i* (left), and for model *M* (right).

**Fig 5.**
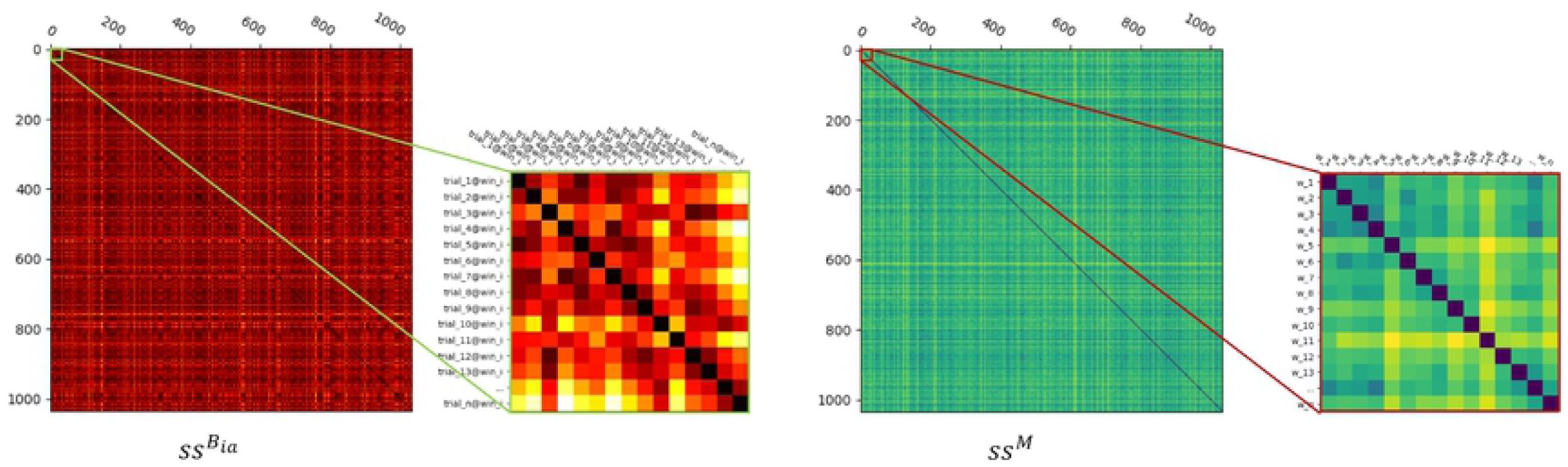
Similarity matrix for brain activity and for model. Similarity matrix for brain activity in region *a* and time *i* (left), and for model *M* (right). The matrices have been zoomed in to show their fine-grained structure and the labels of their rows and columns.

The similarity score is estimated by taking Pearson’s correlation coefficient between the upper off-diagonal triangle of the [*n × n*] symmetric paired similarity matrices (*ss*^*M*^ and 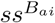) (Fig 6). These scores quantify the extent to which the similarity across stimuli is similarly represented by the model *M* and by brain activity in anatomical region *a* and time *i*. These measures are repeated across time *t* and anatomical regions *r* (Fig 7).

**Fig 6.**
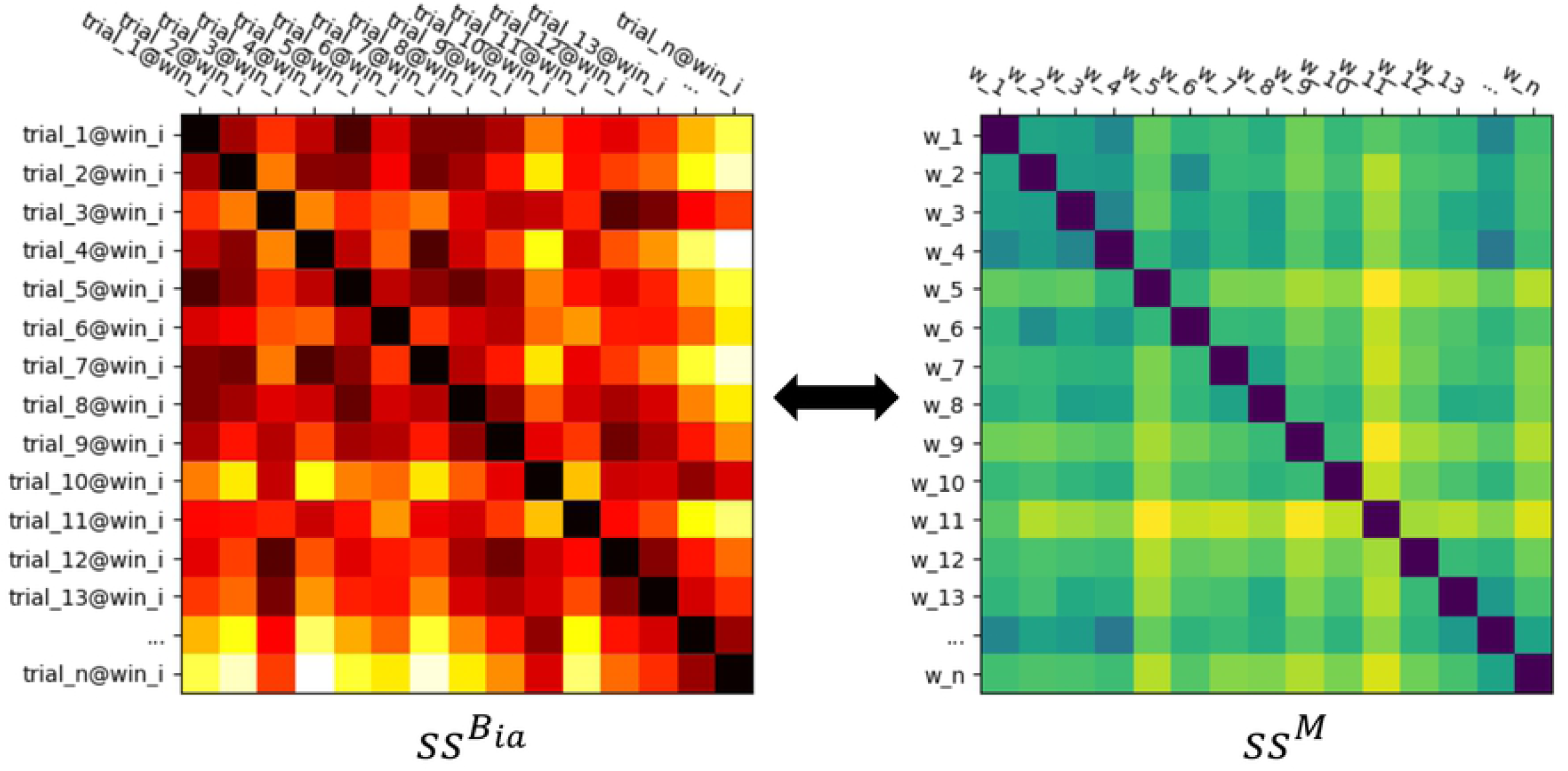
Representational Similarity Analysis (RSA). RSA is conducted by computing the similarity score between the brain space similarity matrices (left) and model similarity matrix (right). Here, the brain similarity matrix consists of the pairwise similarity scores among trials at anatomical region *a* and time window *i*.

**Fig 7.**
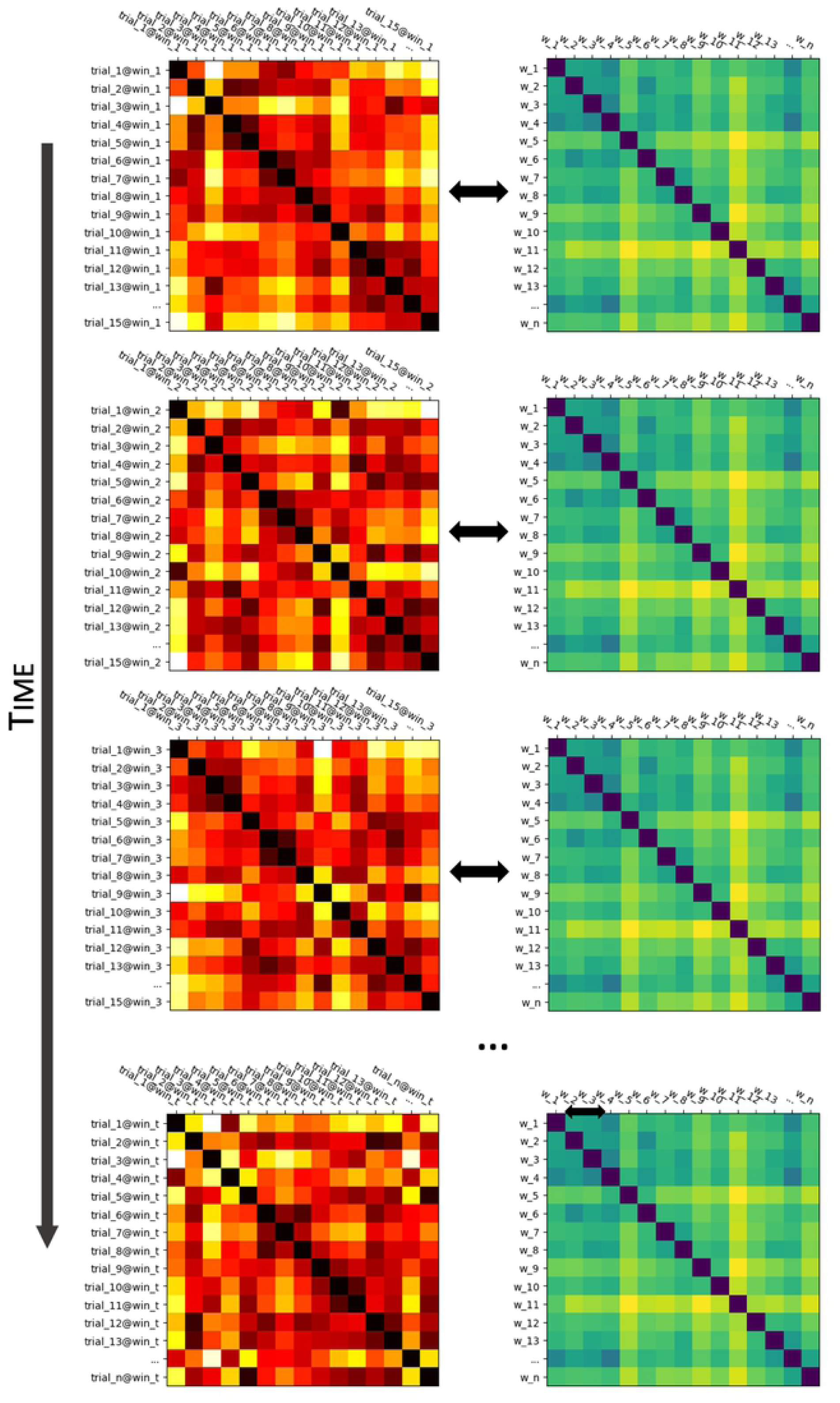
RSA over time. RSA is conducted over time, meaning that a brain space similarity matrix is computed for each separate time window of the MEG trials.

Therefore for each anatomical region, we obtained a representation of similarity between model and brain activity as a function of time from 0 to 500 ms after word onset (in windows of 30 ms, every 16 ms).

The analyses are conducted at the single-subject level up to this stage. The result of the temporal RSA for model *M* thus consists of 74 matrices, one for each subject, of size [*s × u*] where *s* is the number of anatomical regions and *u* the number of temporal windows.

### Group-level analysis

Given 74 per-subject [*s × u*] matrices, we want to obtain a single group-level matrix describing the similarity between model *M* and brain activity across anatomical regions and over time.

Typically, group-level RSA results are obtained by averaging similarity matrices across subjects before computing the similarity score with the model similarity matrix. Cross-subject averaging requires that all matrices have the same size and that the row and columns of the matrices correspond to the same words across matrices. This is not the case for this dataset. First, the subjects are grouped in cohorts of about 17, each of which was presented with a different subset of the 360 stimulus sentences. This means that not all 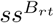 have rows and columns corresponding to the same words. Moreover, the trial selection procedure in the per-subject pre-processing consisted of discarding trials with irreparable artifacts. Therefore, even if two subjects were presented with the same set of sentences, there is a chance that they would have a different set of corresponding MEG trials due to the different occurrence of signal artifacts.

In our analysis, we computed t-statistics over subjects for each region/time combination independently. We computed a one-sided t-score for each of our computational models 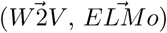. We also computed a one-side t-score between the scores obtained by our aggregate models and word2vec 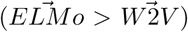. The results of these t-statistics are thresholded at *p<*0.05. Given the exploratory nature of this study, no statistic correction was applied in order not to obliterate the possible small effects detected by the RSA.

## Results

The results of the analyses are split in two main parts. In the first part, we report the results of each of the two embedding models (word2vec and ELMo) separately. In the second part of the analyses, the similarity scores of word2vec and ELMo are contrasted to each other.

Results are provided at the whole-brain level, displaying the model–brain similarities at 5 distinct time points: 150, 250, 350, 450, and 550 milliseconds after word onset.

### Models

In this section, we report the results of the RSA analyses of the two computational models separately.

The embedding model (Fig 8) which does not include contextual information, word2vec, returns lower similarities overall over time when correlated with brain activity, but more so from about 300 ms post word onset. For earlier latencies, word2vec shows significant activity in the left middle and inferior temporal gyri. Significant similarity with brain activity is also observed around 400 ms in the left posterior superior temporal gyrus.

**Fig 8.**
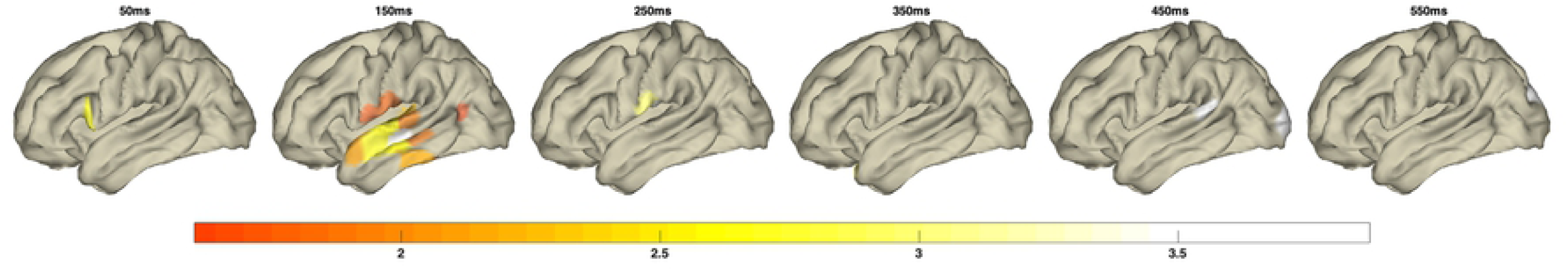
Non-contextualized model (word2vec) This figure displays the results of the RSA between MEG activity and word2vec embeddings (t-values, *p <* 0.05).

The contextualized model (Fig 9), ELMo, instead exhibits an overall significant similarity with brain activity between 300 and 500 ms in the left frontal, prefrontal, and left anterior temporal regions. In particular, ELMo shows significant similarity with the left inferior temporal gyrus and the left anterior temporal cortex around 400 ms.

**Fig 9.**
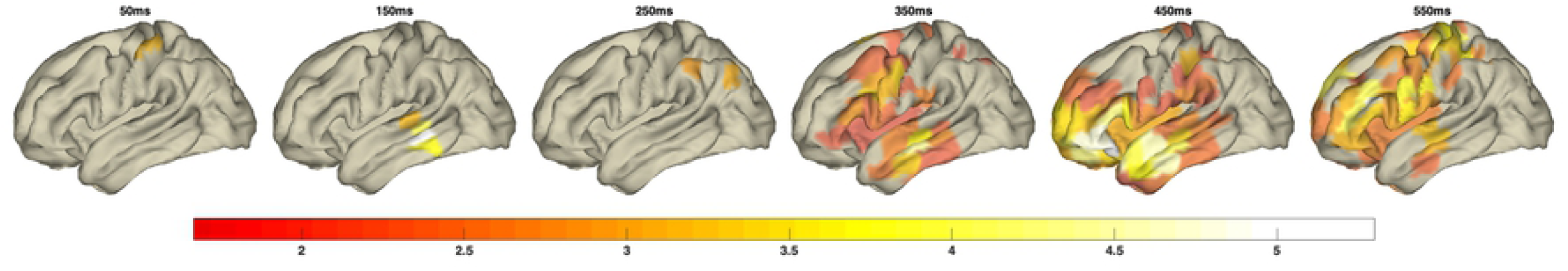
Contextualized model (ELMo) This figure displays the results of the RSA between MEG activity and and ELMo embeddings (t-values, p < 0:05).

### Comparison

In Fig 10 we show the results of a direct comparison between ELMo and word2vec 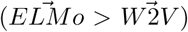.

ELMo shows significantly higher similarity to brain activity as compared to word2vec in the left anterior temporal lobe and in the left inferior frontal gyrus around 400 ms post word onset. The higher scores are also observed for the left middle frontal and left prefrontal regions around 500 ms.

**Fig 10.**
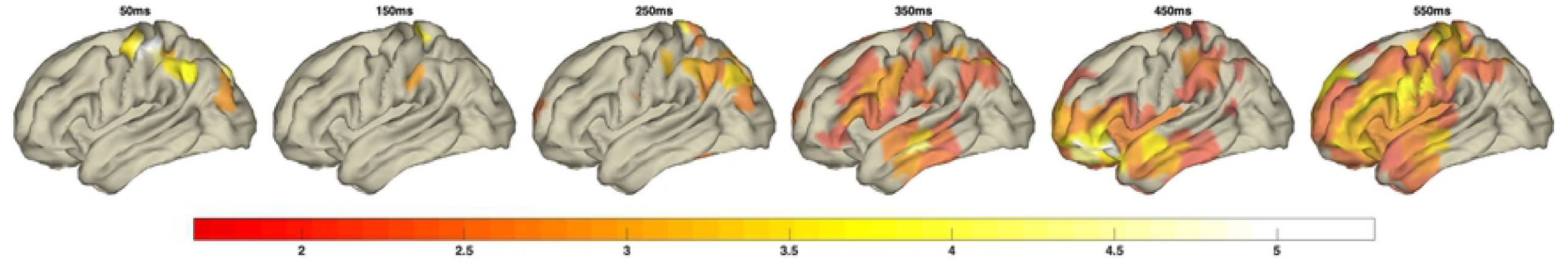
Comparison between contextualized (ELMo) and non-contextualized model (word2vec) This figure displays the comparison the correlations between MEG activity and ELMo and MEG activity and word2vec (t-values, *p <* 0.05).

## Discussion

In Section 4, we observed that contextual and non-contextualized models yield qualitatively different results with regard to the timing and location of their similarity to MEG-recorded brain activity. In this section, we discuss the implications of these findings in light of the nature of the models and of the brain processing of natural language, as introduced in Sections 4 and 4.

We believe that computational word embedding models help in probing the nature of the neural representations correlated to memory retrieval and to integration. This is because they make the distinction between these two phenomena more computationally specific. When discussing the nature of retrieval and integration, Baggio & Hagoort (2011) [7] argue that approaches based on formal semantics might not be realistic models of how the brain implements these two operations. In agreement with Seuren (2009) [47], they state that formal semantics disregards natural language as a psychological phenomenon. They continue stating their desire to develop an account “that adheres to cognitive realism, in that it explains how language users derive meaning and how the human brain instantiates the neural architecture necessary to achieve this feat”. We believe that distributional semantic models, of which contextualized embeddings are the most advanced version, have already proven their cognitive realism by being good models of human behavioral – e.g. semantic similarity judgment [48] – and neural [49, 50] data. Moreover, at the dawn of the field, distributional models – e.g., Latent Semantic Analysis [51] – were actually developed as cognitive models to answer questions on how children acquire word meaning, and how humans react to semantic similarity and relatedness. In light of the above considerations, we think that the models presented in the study might offer a cognitively realistic approximation of what is going on in the brain during memory retrieval and integration.

In the remainder of this section, we will discuss the effect of contextualization on the similarities between computational representations and brain activity (Section 4). We will specifically focus on the implications of these findings regarding the role of the anterior temporal lobe 4 and of activity peaking around 400 ms after stimulus onset (Section 4). We will also discuss the plausibility of the models chosen for the present study (Section 4).

### The effect of contextualization on model-brain similarity

ELMo creates vectorial representation of words (embeddings) dependent on the context in which those words occur (Section 4). This contrasts with the nature of embeddings generated by a model such as word2vec (Section 4) that, once trained, always generates the same embeddings for a word regardless of its context.

Contextualized word embeddings generated by ELMo display higher similarity with brain activity in the frontal and anterior temporal regions of the brain around 300 and 400 ms post word onset; in other words, in areas and in a time frame compatible with integration processes. Integration refers to the integration of a linguistic unit (a word for instance) in the context provided by the other linguistic input in which it happens to be contained.

Non-contextualized word embeddings generated by word2vec exhibit a somewhat opposite behavior, showing higher similarity with earlier (100–200 ms) activity in lateral temporal regions. These regions are supposed to implement long-term memory (semantic memory) retrieval.

The fact that ELMo and word2vec embeddings display such different behaviors with regard to brain activity can be reconciled with the division of labor predicated by models such as the MUC model [28], especially concerning the distinction between retrieval from memory and contextual integration.

### The role of the anterior temporal lobe in integration

The fact that contextualized embeddings show higher similarity in the left anterior temporal lobe might indicate that this region plays a role in integration. This is in line with several experimental studies reporting the involvement of this area in lexical, semantic and syntactic integration. Works such as Mazoyer & al. (1993) [29], Stowe & al. (1998) [30], Friederici & al. (2000) [15], Humphries & al. (2006) [31], and Humphries & al. (2007) [32] arrived at the conclusion that the anterior temporal lobe is somehow involved in sentence-level integration, after observing an increase in activity in this area during the presentation of sentences as compared to word lists. In addition, a series of other studies have confirmed the role of ATP in processing composition by showing that its activity is modulated by the type of syntactic relation holding between two words being integrated [33–36, 36, 37].

The left ATP is also considered central in semantic memory. This was confirmed by studies on patients suffering from semantic dementia [21, 22, 52, 53], and several functional neuroimaging studies [23–26, 54]. These findings are summarized by Patterson & al. 2007 [27], who hypothesized that concepts are represented by a network of sensorimotor representations united by the left ATP.

Contextualized models do not show high correlations in the ATP at latencies that are associated with memory-related processing: 100–200 ms after word presentation. The results, therefore, seem to indicate that contextualized embeddings approximate representations that have to do with the integration into context and not with lexical retrieval from memory per se. If this were the case, we would have expected similar correlations in the ATP both at memory retrieval (100–200 ms) and integration latencies (400 ms). Conversely, a tentative interpretation with regard to the role of ATP could be that activity in this region is different when semantic memory operations and contextual integration processes are carried out.

### The role of N400 in integration

The results presented in Section 4 do confirm not only the anatomical loci of memory and integration, but also provide indirect suggestions on the role of the N400 [41]. It has been debated in the literature whether the N400 is best characterized as playing a role in combinatorial (integration) or non-combinatorial (retrieval from memory) processes. Baggio & Hagoort (2011) [7] and Hagoort & al. (2008) [40] refrain from providing a stance on the matter because of the difficulty of devising convincing task-dependent experimental designs that are able to disentangle combinatorial and non-combinatorial semantic processes. Here again, we believe that the present computational modeling approach may provide an answer to the question. The results seem to point more towards a combinatorial process for the N400, given that the contextualized model maps to brain activity corresponding to its latencies. It is an indirect proof, but a proof pointing in this direction nonetheless. These results are in line with studies that link the role of the N400 to context processing [38, 39].

### The plausibility of bi-directional RNNs

ELMo is essentially a bi-directional recurrent neural network-based language model that integrates a word with its preceding and following context. Bi-directional recurrent language models seem to violate the assumption that human language processing proceeds left-to-right and word-by-word. Although this is trivially true for listening, it is worth noting that studies of reading behavior tend to describe a more nuanced situation. It is consistently reported in the eye-tracking of reading literature that between 15 and 20 % of eye movements proceed right-to-left (regressions) for left-to-right languages such as English or Dutch, and the opposite occurs for right-to-left languages such as Arabic [55]. Moreover, a number of “jump-ahead” eye movements are also commonly observed, indicating that humans either skip information that is deemed irrelevant for the processing of a linguistic item or that they look ahead in order to collect contextual information to the right of a word. This indicates that the preceding linguistic sequence is not always the only contextual material employed in the processing and interpretation of a word or of a sentence as a whole.

## Conclusions

Recent developments in computational linguistics have created a new family of models which generate word embeddings that compute their representations with information derived from the context in which words are used. In this study, we have adopted one particular contextualized model, ELMo, as an approximation of the result of integration processes in the human brain during natural language comprehension. We contrasted ELMo with word2vec, a non-contextual embeddings model. Starting from the distinction between semantic memory retrieval (implemented in left temporal regions and activated around 200 ms after the onset of an incoming word) and word integration into context (carried out in left inferior frontal and left anterior temporal regions around 400 ms after word onset), we observed that non-contextualized models correlate with activity only in regions and latencies associated with semantic memory. In contrast, contextualized models correlate with activity in areas and latencies associated with word integration in context. These results confirm the functional and physiological distinction between memory and integration. Moreover, they provide some insight into the role of the left IFG, an area involved in integration and whose activity might temporarily store contextualized lexical representations. The results also point towards an involvement of the left anterior temporal lobe in integration, an area that was already linked to semantic combinatorial processes and which nonetheless received less attention in the theories of the architecture of the language system adopted in this study.

By highlighting a parallelism between models and brain activity, our results offer a contribution to the understanding of the division of labor at the cortical level between areas encoding lexical items in isolation and areas sensitive to the use of those items in context.

## Bibliography

## Footnotes

1 https://github.com/HIT-SCIR/ELMoForManyLangs [43, 44]

